# Humans can track but fail to predict accelerating objects

**DOI:** 10.1101/2021.11.20.469397

**Authors:** Philipp Kreyenmeier, Luca Kämmer, Jolande Fooken, Miriam Spering

## Abstract

Objects in our visual environment often move unpredictably and can suddenly speed up or slow down. The ability to account for acceleration when interacting with moving objects can be critical for survival. Here, we investigate how human observers track an accelerating target with their eyes and predict its time of reappearance after a temporal occlusion by making an interceptive hand movement. Before occlusion, the target was initially visible and accelerated for a brief period. We tested how observers integrated target motion information by comparing three alternative models that predicted time-to-contact (TTC) based on the (1) final target velocity sample before occlusion, (2) average target velocity before occlusion, or (3) target acceleration. We show that visually-guided smooth pursuit eye movements reliably reflect target acceleration prior to occlusion. However, systematic saccade and manual interception timing errors reveal an inability to consider acceleration when predicting TTC. Interception timing is best described by the final velocity model that relies on extrapolating the last available velocity sample before occlusion. These findings provide compelling evidence for differential acceleration integration mechanisms in vision-guided eye movements and prediction-guided interception and a mechanistic explanation for the function and failure of interactions with accelerating objects.

## Introduction

Seeing and perceiving object motion is a vital capability of the primate visual system. Animals hunting for prey or pedestrians crossing a street must be able to act upon a target’s speed, direction, and sudden target accelerations or decelerations. Even though acceleration is an important feature that describes the behavior of many moving objects, it is well established that the primate perceptual system is largely insensitive to it (Gottsdanker et al., 1961; Calderone & Kaiser, 1989; Werkhoven et al., 1992). How we incorporate visual acceleration signals into motor commands to interact with moving objects is still not fully understood and would require a systematic comparison of different effector types. Here, we evaluate human observers’ ability to track accelerating targets with their eyes and to predict accelerating target trajectories for manual target interception.

Tracking visual object motion with the eyes and predicting an object’s motion path are two fundamental abilities that both rely on decoding visual motion (Fiehler et al., 2019) and can inform consecutive hand movements to catch, hit or otherwise intercept the target (Mrotek & Soechting, 2007; Mrotek, 2013; Diaz et al., 2013; Cesqui et al., 2015; Fooken et al., 2016; de la Malla et al., 2017; Goettker et al., 2019; De Brouwer et al., 2021; Fooken et al., 2021). However, tracking and predicting differ with regard to the sensory signals that drive them. Whereas tracking relies on combined visual and nonvisual signals (e.g., feedback from motor command units; Lisberger, 2015), predicting a trajectory requires memory of previously seen motion (Orban de Xivry et al., 2013; Kowler et al., 2019; Rust & Palmer, 2021). Whether and how acceleration signals are used for tracking and predicting is the focus of this study.

The surprisingly limited ability to perceptually detect and evaluate accelerating visual targets in humans (Gottsdanker et al., 1961; Calderone & Kaiser, 1989; Werkhoven et al., 1992; Brouwer et al., 2002; Watamaniuk & Heinen, 2003; Benguigui et al., 2003; Mueller et al., 2017), led researchers to conclude that rather than perceiving acceleration directly, humans might merely detect changes in velocity and derive acceleration signals indirectly (Gottsdanker et al., 1961). This view is congruent with findings obtained in neurophysiological studies in macaque monkeys. Single neurons in motion-sensitive extrastriate cortex (area MT) are not tuned to acceleration. Instead, acceleration signals can only be indirectly decoded from populations of speed-sensitive neurons (Lisberger & Movshon, 1999; Price et al., 2005).

By contrast, human eye movements appear to be sensitive to visual acceleration. Smooth pursuit—slow rotations of the eyes that allow continuous tracking of moving objects—can match target velocity perturbations (Tavassoli & Ringach, 2010; Brostek et al., 2017). When tracking an accelerating target throughout a brief occlusion period, eye movements (pursuit and saccades) scale with object acceleration in anticipation of target reappearance (Bennett & Barnes, 2006; Bennett et al., 2007; Bennett & Benguigui, 2013). These findings indicate that the pursuit system can extract target acceleration effectively and use this signal to predictively drive continuous tracking eye movements.

Yet, when observers are asked to judge the time or position of target reappearance or time-to-contact (TTC) by making an interceptive hand movement or press a button, systematic errors indicate an inability to consider target acceleration for action-related motion prediction (Benguigui et al., 2003; Benguigui & Bennett, 2010; Bennett & Benguigui, 2013; Brenner & Smeets, 2015; Brenner et al., 2016; Brouwer et al., 2002; Port et al., 1997; Dubrowski & Carnahan, 2002). Observers fail to adjust the timing of their hand movement to the target’s acceleration, resulting in interceptions that are too early and ahead of the target when it decelerates, and interceptions that are too late and behind the target when it accelerates.

In sum, whereas visually-guided eye movements to accelerating targets appear to be relatively responsive to acceleration, hand movements to accelerating targets are prone to systematic errors that indicate inability to account for acceleration. To investigate whether acceleration signals are used differently when tracking or predicting a visual target, we compare visually-guided eye movements and interceptive eye and hand movements directly. In a track-intercept task, observers were asked to track an accelerating target with their eyes before it was temporarily occluded, and to intercept it immediately upon reappearance with a quick pointing movement (**Fig. 1A,B**). This task requires observers to accurately decode target acceleration during the initial target presentation, to predict the TTC, and to adjust the execution of their manual interception in order to hit the target at the right time.

**Figure 1.**
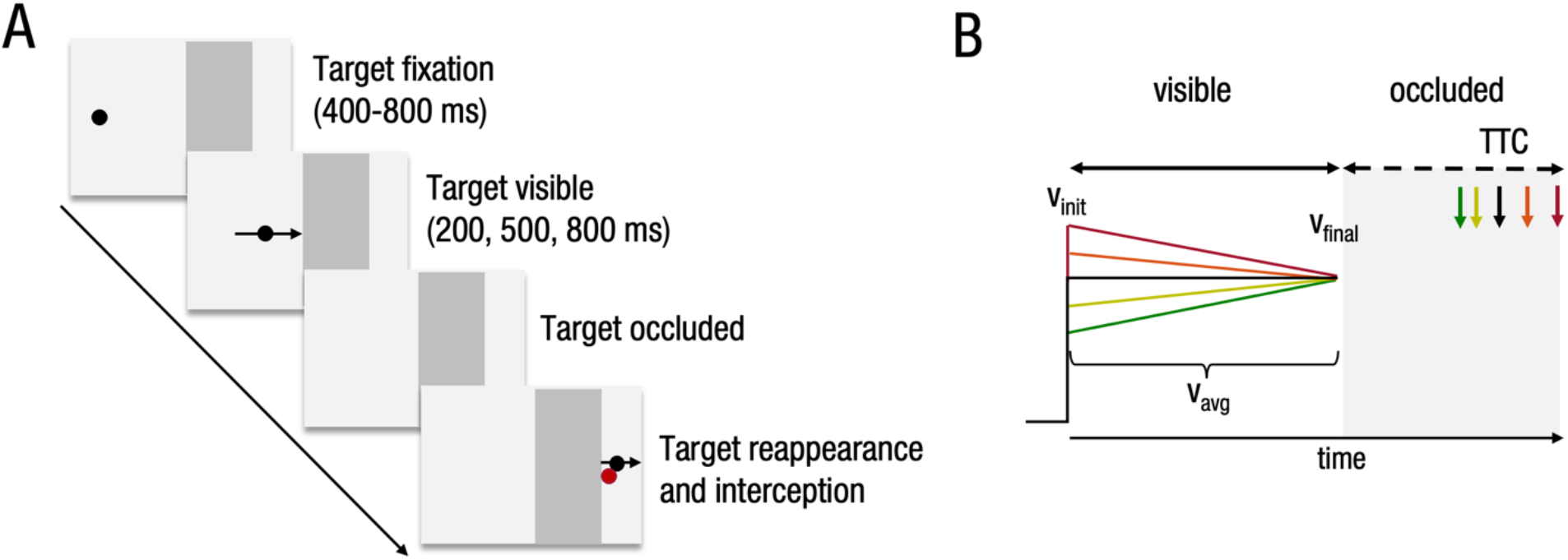
**(A)** Task procedure. A black disc moved across a grey monitor background from left to right at a constant rate of acceleration. After an initial period during which the target was visible, it moved behind an occluder of a fixed width (13.4°) and then reappeared. Participants had to estimate the time of reappearance (equivalent to time-to-contact; TTC) and intercept the target with a rapid pointing movement of their right index finger (red dot). **(B)** Target parameters. Targets moved with a variable initial velocity (v_init_) and accelerated or decelerated at a constant rate. The initial and average velocities (vavg) of the targets were related in such a way with acceleration rate that all targets reached the occluder with the same velocity (v_final_) of 20°/s.

Importantly, our task design allows us to empirically test and compare the predictions of three competing models of how target acceleration signals are integrated with velocity signals (Bennett et al., 2007; Heinen, 2007) and utilized to predict TTC. Our data reveal that tracking (visually-guided smooth pursuit eye movements) accurately distinguish between accelerating target trajectories. However, observers fail to account for acceleration when predicting target motion for ocular (saccadic) and manual interception. These interception findings confirm predictions of the final velocity model, which postulates that observers continuously update target velocity and predict target reappearance based on the last available velocity sample. They are less compatible with two alternative accounts for acceleration signal integration: the average velocity model, which assumes that observers base their interception timing on the average target velocity, or the acceleration model, suggesting that observers consider target acceleration for interception at least to some degree. Taken together, our results imply that the prediction of accelerating targets for interceptive eye and hand movements does not benefit from the pursuit system’s ability to accurately track accelerating targets. Our findings reveal a dissociation between visually-guided pursuit and interception, and at the same time a striking parallel in how both effectors—eye and hand—are limited in utilizing target acceleration for prediction-based interception.

## Results

Observers (*n* = 16) performed a track-intercept task in which they viewed a moving disc that disappeared behind an occluder after a presentation time of 200, 500, or 800 ms and then reappeared for 100 ms (**Fig. 1A**). We instructed observers to intercept the target at the estimated time of reappearance (equivalent to TTC) with a quick pointing movement while recording their eye and hand movements (see **Materials and Methods**). In each trial, the target moved along a horizontal linear path either at a constant velocity (no acceleration, 0°/s^2^: control condition) or at a constant rate of velocity change (deceleration: −8 or −4°/s^2^; acceleration: +4 or +8°/s^2^). Initial target velocity depended on the rate of change in velocity and was matched so that all targets reached the occluder with the same final velocity of 20°/s (**Fig. 1B**).

We analysed the data in four parts: First, we investigated whether target acceleration was reflected in visually-guided smooth pursuit eye movements during initial target presentation. Second, we assessed the effect of target acceleration on the timing of the interceptive hand movement. We extended this analysis to interceptive saccades, which were made instinctively by all observers and coincided with the hand movement. In the third part, we compared the performance of three acceleration integration models for the prediction of TTC in observers’ eye and hand movement data. In the first three parts, we focused on the longest presentation duration (800 ms) because it yielded the most reliable eye movement responses. In the fourth part, we investigated eye and hand movements in task conditions, in which observers’ ability to accurately track and decode target acceleration was restricted by shorter presentation durations (200 or 500 ms).

### Target Acceleration is Accurately Reflected in Smooth Pursuit

To investigate whether smooth pursuit is sensitive to acceleration signals, we analysed observers’ eye movements while they tracked the visual target before occlusion. Two example trials, in which the target either accelerated (**Fig. 2A**) or decelerated (**Fig. 2B**) show that observers tracked with a combination of smooth pursuit and saccadic eye movements during the initial target presentation. After pursuit initiation, an initial catch-up saccade aligned the eyes with the target and was typically followed by a period of closed-loop smooth pursuit, during which eye velocity closely matched the continuously changing target velocity. In some trials, smooth pursuit was supported by additional catch-up saccades (**Fig. 2B**). Around the time of target occlusion, observers reliably made a distinctly identifiable targeting saccade of relatively large amplitude to the end of the occluder, where they intercepted the target with a pointing movement of their right index finger (see 3D hand trajectory from a single trial in **Fig. 2C**).

**Figure 2.**
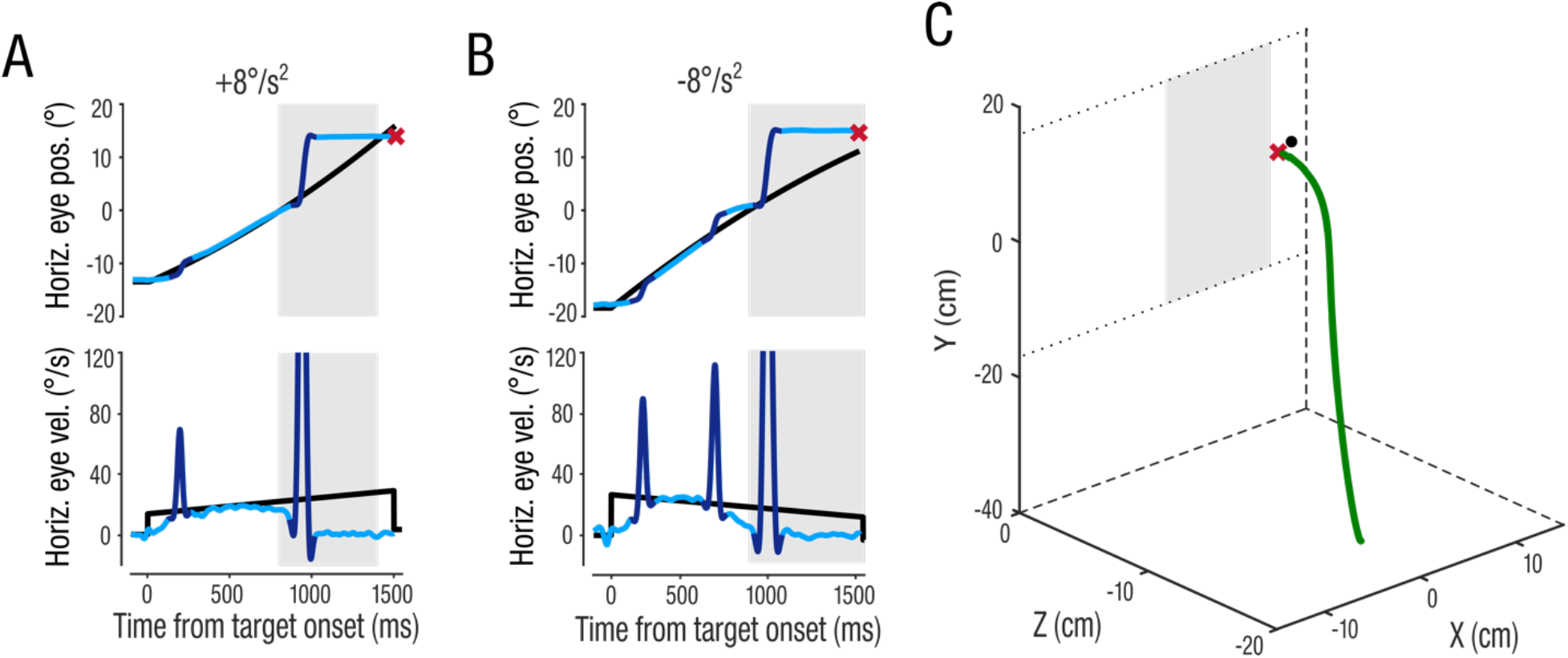
Single trial eye and hand movements from one representative observer. (**A** and **B**) show two trials with a +8°/s^2^ accelerating target (**A**) or −8°/s^2^ decelerating target (**B**). Light blue traces indicate smooth pursuit components, dark blue traces represent saccades. Upper panels show the horizontal position of the eye (blue) and target (black) locked to target motion onset. The red ‘x’ represents the interception position and time. Lower panels show horizontal velocity of the eyes and target over time. Grey area represents the time of target occlusion. (**C**) shows the 3D-hand position trace (green) from the same trial as in (**A**). The 2D interception position on the screen is indicated by the red ‘x’ and the target position at the time of interception in represented by the black disc. The grey area illustrates the position of the occluder on the screen. Dotted lines in the x-y plane illustrate the upper and bottom edges of the screen.

Average pursuit velocity traces across trials and observers reflect the velocity profiles of the different accelerating targets (**Fig. 3A**). To account for imperfect velocity gain and individual differences, we normalized eye velocity by subtracting average eye velocity in the control condition (0°/s^2^) from average eye velocity in acceleration conditions for each observer. Normalized eye velocity profiles (**Fig. 3B**) reveal a clear signature of target acceleration and show that observers were able to continuously match their eyes to the changing target velocities over time. These observations were confirmed by a main effect of *acceleration* on closed-loop eye acceleration (*F*(3,45) = 11.3; *p* < .001; *η_p_^2^* = .44; **Fig. 3C**), analysed between offset of the first catch-up saccade and onset of target occlusion. Because eye acceleration is the second derivative of eye position and therefore a noisier eye movement metric, we confirmed these results by additionally analysing eye velocity at two single time points: at the beginning of the closed-loop pursuit phase (after the initial catch-up saccade; approx. 320 ms after target onset) and before occlusion, when target velocity was identical across conditions. Pursuit velocity accurately reflected target velocities at both time points, indicated by a main effect of target *acceleration* at the beginning of the closed-loop phase (one-way rmANOVA: *F*(3,45) = 82.0; *p* < .001; *η_p_^2^* = .85; **Fig. 3D**) and similar pursuit velocities across acceleration levels before occlusion (*F*(3,45) = 1.83; *p* = .18; *η_p_^2^* = .11; **Fig. 3E**). Note, a lack of main effect at the latter time point would be expected if the eyes matched the identical target velocities at the time of occlusion. Thus, the observed changes in smooth pursuit acceleration and velocity scaled with changes in target velocity across the entire time period in which the target was visible. Together, these results suggest that observers were able to accurately extract acceleration information, implying that acceleration signals are available to the oculomotor system to be utilized for target prediction and interception in this system, and potentially in the hand motor system as well.

**Figure 3.**
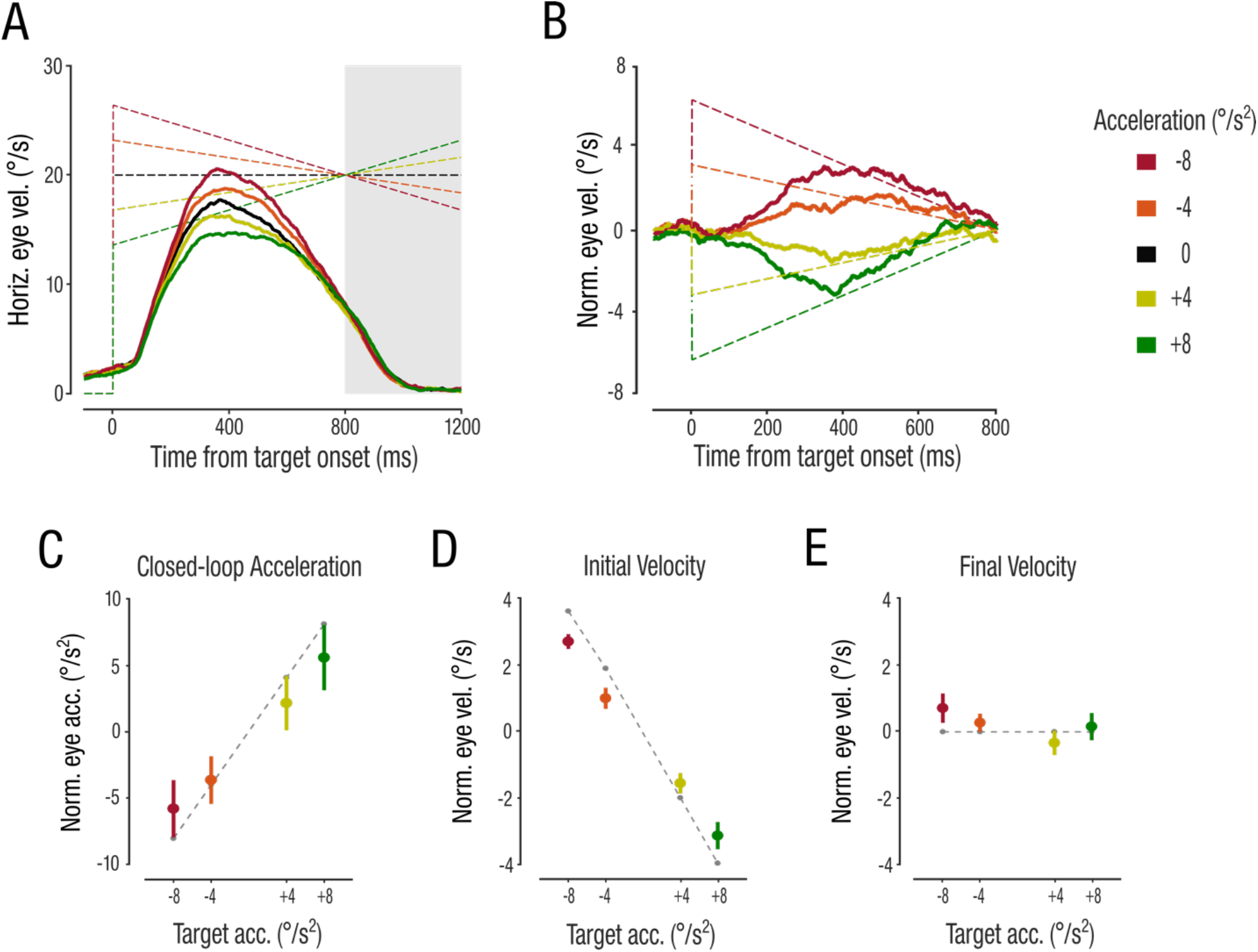
Average de-saccaded smooth pursuit data. (**A**) Horizontal eye velocity traces over time, aligned to target onset. Dashed lines represent target velocity profiles. Grey area indicates the time of target occlusion. (**B**) Normalized pursuit velocity after subtraction of the control condition. (**C**) Normalized, closed-loop smooth pursuit acceleration averaged over the time from the beginning of the closed-loop pursuit phase (defined as the time of the initial catch-up saccade offset) until occlusion onset. Grey dashed line shows rates of target acceleration. **(D**) Normalized pursuit velocities at the beginning of the closed-loop pursuit phase. Grey dashed line shows the mean target velocities at the time of the initial catch-up saccade offset. Note that accelerating and decelerating targets moved with maximally different velocities at the beginning of the trajectory in order to reach the occluder with the exact same velocity of 20°/s, resulting in high initial velocity for decelerating and low velocity for accelerating targets, explaining the negative slope in this data panel. (**E**) Normalized pursuit velocity at the time of target occlusion. Grey dashed line shows the target velocities at the time of occlusion. All data represent the mean across observers ± 1 standard error of the mean (SEM).

### Target Acceleration Causes Systematic Interception Errors

Consequently, we next asked whether target acceleration was reflected in manual interception. We analyzed the constant error of interception, i.e., the temporal error describing by how much a target was hit too early or late. We also analyzed hand movement kinematics (hand latency, movement time, peak hand velocity, and interception time). If hand movement data reflect acceleration akin to what we observed in pursuit eye movements, we would expect observers to start moving their hand sooner and intercept earlier for accelerating targets and intercept later for decelerating targets, relative to the zero-acceleration control condition. An acceleration-adjustment of the hand movement would thus lead to accurate interceptions, whereas insufficient adjustment would lead to systematic constant errors.

As instructed, observers intercepted the target close to the occluder’s right edge (mean distance 1.2 cm ± .48 cm; **Fig. 4A**). They initiated the hand movement at a latency of 216 ms (± 136 ms) relative to occlusion onset and the hand movement took 474 ms (± 118 ms) to complete. In contrast to what we would expect if acceleration signals directly impacted interception, observers hit accelerating targets too late (i.e., the target had already passed the interception position) and decelerating targets too early (i.e., the target had not yet reached the interception position), confirmed by a main effect of target acceleration on constant interception error (*F*(4,60) = 127.3; *p* < .001; *η_p_^2^* = .92; **Fig. 4B**). This systematic acceleration effect on the constant error suggests that observers failed to consider target acceleration to correctly estimate TTC—a target whose acceleration is not decoded would always be intercepted too late, and a decelerating target would always be intercepted too early. Interestingly—and in contrast to what we would expect, if observers considered target acceleration—observers began their hand movement later in response to target acceleration and earlier for deceleration (main effect of *acceleration*: *F*(4,60) = 6.18; *p* = .009; *η_p_^2^* = .29; **Fig. 4C**). We did not find any evidence that observers corrected this bias mid-flight by adjusting the speed of their interception movement (no main effect of *acceleration* on movement time or peak hand velocity, all *F*(4,60) < 1.31; *p* > .28). Consequently, the time of manual interception followed the same pattern as the hand latency—later for accelerating targets and earlier for decelerating targets (*F*(4,60) = 6.20; *p* = .01; *η_p_^2^* = .29; **Fig. 4D**).

**Figure 4.**
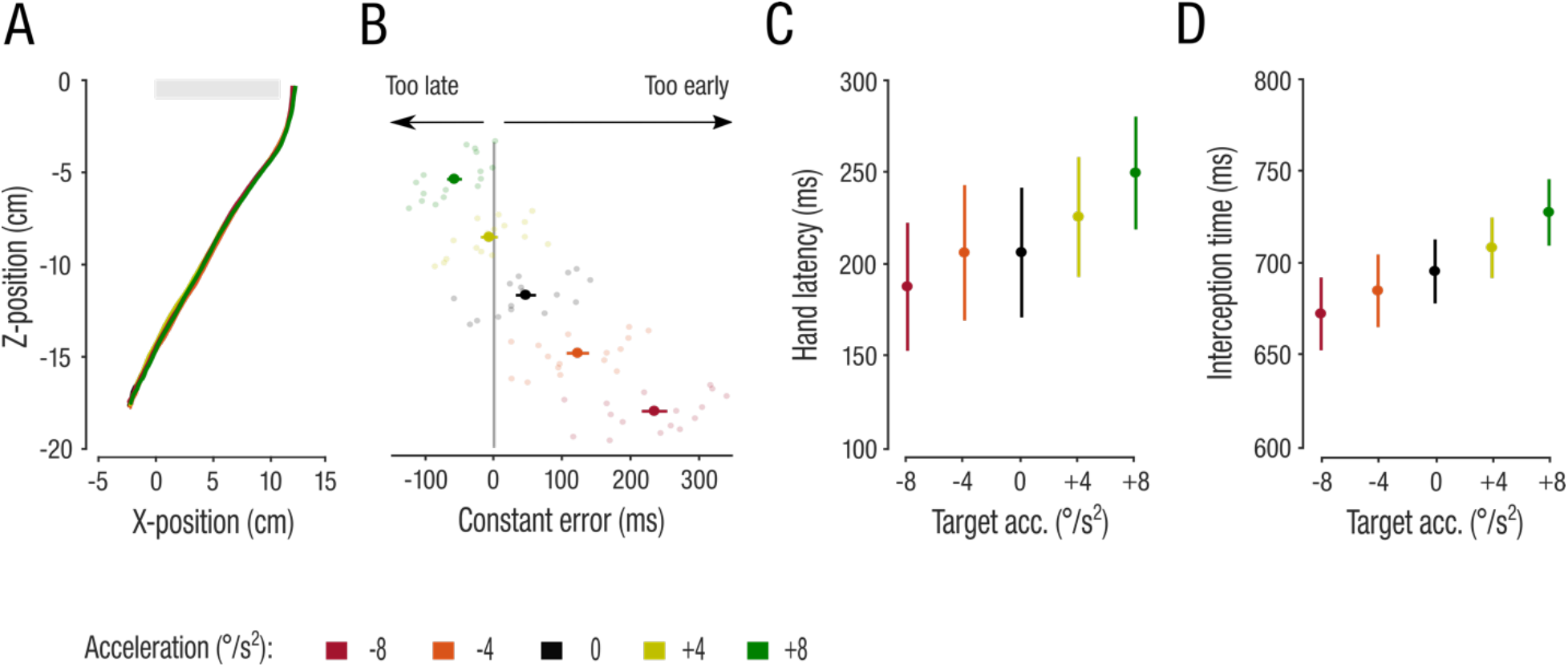
Hand movement kinematics. (**A**) Top view of the mean hand movement paths in x- and z-coordinates. The grey bar illustrates the x-position of the occluder on the screen. (**B**) Mean and individual interception constant errors. Semi-transparent dots represent individual observers’ mean performances. Negative values indicate interceptions that occurred after the target reached the end of the occluder (too late) and positive values indicate interceptions occurring before the target reached the end of the occluder (too early). (**C**) Mean hand latencies measured from target occlusion onset. (**D**) Mean interception times measured from target occlusion onset. Conventions are the same as in Fig. 3.

### Which Target Features Determine TTC Predictions?

The observed constant interception errors and the systematic biases in hand movement kinematics (later interception for accelerating targets and earlier interception for decelerating targets) suggest that observers were not able to correctly adjust their interception movement according to target acceleration. In contrast, observers were able to accurately track accelerating targets until occlusion. However, we do not know whether the oculomotor system uses the acceleration readout during visually-driven pursuit to inform predictive eye movements. In our paradigm, observers typically stopped pursuing the target smoothly and shifted their eyes from the occluder’s left edge to the right edge by making predictive saccades to the point of reappearance (**Fig. 2A,B**). We next asked whether these predictive eye movements resembled the acceleration-sensitive pattern observed in pursuit or were similarly biased compared to interceptive hand movements. We analysed the landing time of the eye at the interception location as the eye’s interceptive response (TTC_eye_). Observers typically initiated saccades toward the interception location shortly after target occlusion (96 ms ± 81; **Fig. 2A,B**), reaching the end of the occluder approximately 264 ms (± 82 ms) after occlusion onset and 434 ms (± 108 ms) before the time of manual interception. The eyes landed at the interception location later for accelerating targets and earlier for decelerating targets (*F*(4,60) = 4.02; *p* = .039; *η_p_^2^* = .21). Taken together, these findings show that predictive saccades did not follow the same pattern of results as visually-driven pursuit but instead resemble the systematic biases observed in interception.

To formally describe the observed eye and hand interception patterns, we compared the biases in TTC_eye_ and TTC_hand_ to three competing model predictions on how observers might estimate the TTC of accelerating targets (Bennett et al., 2007; Heinen, 2007). As one possibility, observers could continuously update target velocity and estimate TTC based on the last available target velocity sample before occlusion (final velocity model; **Fig. 5A**). Alternatively, observers might use the average velocity during the visible period to estimate target reappearance (average velocity model; **Fig. 5B**). Finally, if observers indeed considered target acceleration, we would predict accurate scaling of the estimated TTC with target acceleration (acceleration model; **Fig. 5C**). We quantified the performance of each model by calculating the root-mean-square error (RMSE) for each observer.

**Figure 5.**
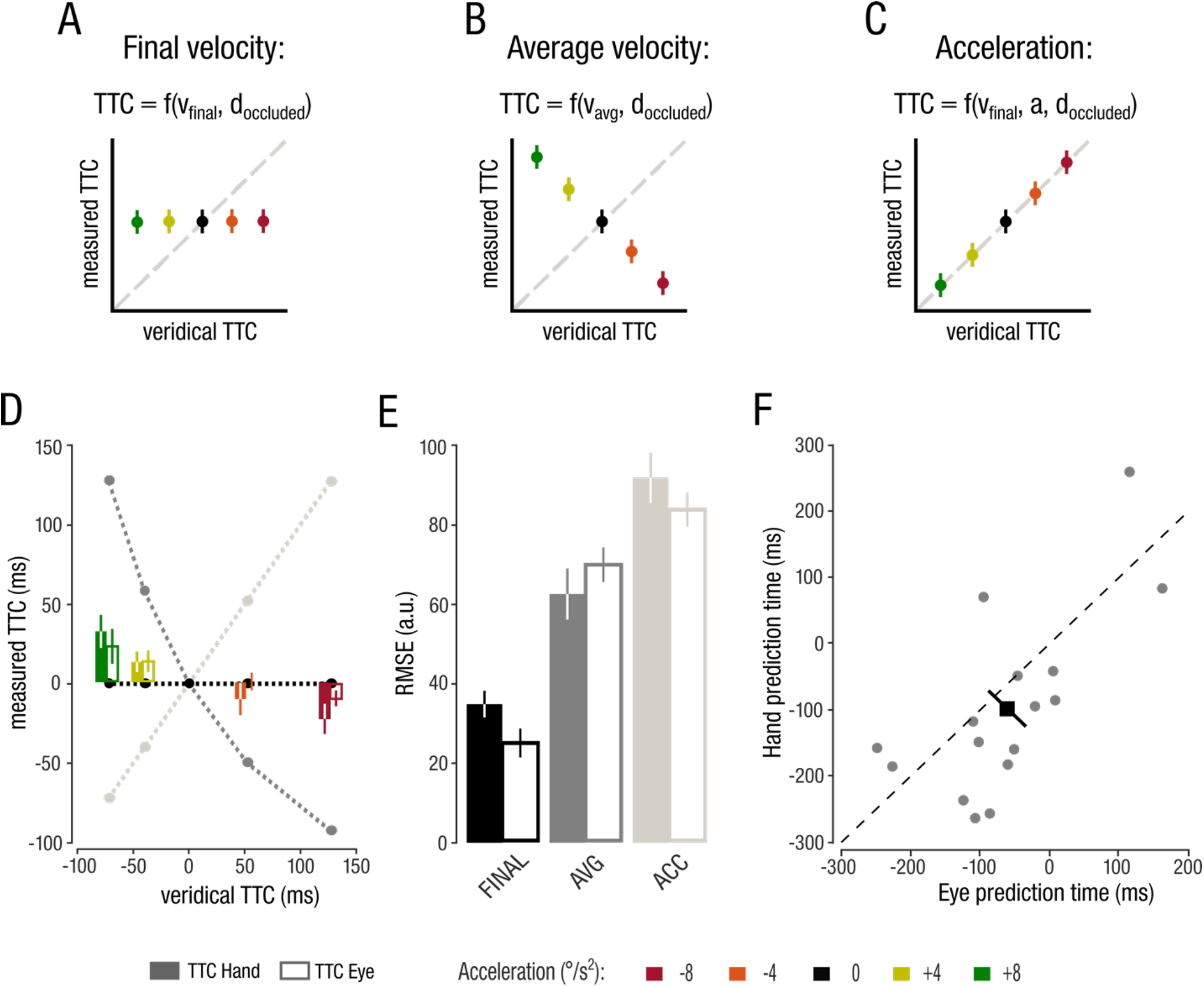
Model comparison. (**A-C**) Three competing models of how observers might predict TTC for interceptive eye and hand movements: (**A**) The final velocity model postulates that observers predict TTC based on v_final_ (identical for all targets in our design, hence predicting a fixed TTC). (**B)** The average velocity model predicts interception timing based on the average target velocity before occlusion, yielding a negative correlation between veridical and measured TTC. (**C**) The acceleration model suggests that observers use target acceleration for interception and predicts the veridical TTC. (**D**) Comparison of model predictions and normalized TTC_eye_ and TTC_hand_ data. (**E**) Root mean squared errors for the competing models on TTC_eye_ and TTC_hand_. Error bars denote ± 1 SEM. (**F**) Scatterplot showing individual eye and hand prediction times. Dashed line shows the line of unity. Black square shows the mean across observers and diagonal errorbars denote the 95% confidence interval (CI) of the mean difference between the eye and hand prediction times.

Using the acceleration model to predict TTC data in eye and hand interception confirms that acceleration is not taken into account (**Fig. 5D**; bright grey line). Although the average velocity model captures the small, reversed trend we observed in TTC_eye_ and TTC_hand_ (**Fig. 5D**; dark grey line, see also **Fig. 4D**), this model also performs poorly at predicting the measured TTC. The final velocity model produced the lowest RMSE for both TTC_eye_ and TTC_hand_ (**Fig. 5D**; black line). A main effect of *model* (*F*(2,30) = 32.94; *p* < .001; *η_p_^2^* = .69) in a 3 (*model*) x 2 (*effector*) rmANOVA on the RMSEs supported these observations (**Fig. 5E**). Post-hoc pairwise comparisons confirmed that the final velocity model produced significantly lower RMSEs compared to the acceleration (TTC_eye_: *t*(15) = 15.55; *p* < .001; *d* = 3.89; TTC_hand_: *t*(15) = 9.91; *p* < .001; *d* = 2.48) and average velocity models (TTC_eye_: *t*(15) = 6.81; *p* < .001; *d* = 1.77; TTC_hand_: *t*(15) = 3.48; *p* = .01; *d* = .87). There was no main effect of *effector* (*F*(1,15) = 2.78; *p* = .12; *η_p_^2^* = .16) and no *model* × *effector* interaction (*F*(2,30) = 3.58; *p* = .06; *η_p_^2^* = .19), indicating that the models performed similarly for TTC_eye_ and TTC_hand_.

The final velocity model predicts that observers rely on the last available velocity sample. To pinpoint the approximate sample observers relied on when estimating TTC, we determined the timepoint along each target’s velocity trajectory that best accounted for each observer’s bias in TTC_eye_ and TTC_hand_. We termed this the eye prediction time or hand prediction time (**Fig. 5F**). A negative value indicates that the observer based the TTC prediction on a velocity sample before occlusion, whereas a positive value would indicate that the TTC prediction was based on a partial extrapolation of the veridical target trajectory during occlusion. We found an average eye prediction time of −62 ms and a hand prediction time of −98 ms (**Fig. 5E**). Both prediction times were significantly different from 0 (TTC_eye_: *t*(15) = −2.39; *p* = .03; *d* = .60; TTC_hand_: *t*(15) = −2.82; *p* = .01; *d* = .70), indicating that observers’ TTC estimates indeed relied on velocity samples before the start of target occlusion. Moreover, prediction times were comparable for eye and hand (*t*(15) = 1.44; *p* = .17; *d* = .30; **Fig. 5F**), indicating that estimates in both effectors relied on similar target velocity signals.

The similarity across effectors poses the question whether TTC_eye_ and TTC_hand_ were correlated on a trial-by-trial basis, which would imply a similarity in the trial-by-trial variability between eye and hand movements. Typically, trial-by-trial correlations are interpreted as evidence for common information sources in the signals that drive eye and hand movements (Sailer et al., 2000). We calculated trial-by-trial correlations between TTC_eye_ and TTC_hand_ for each observer individually and across acceleration conditions. On average, correlations were moderately strong (mean *r* = .41 ± .17; **Fig. 6A**) and significantly different from 0 (*t*(15) = 9.73; *p* < .001; *d* = 2.43). To rule out that the trial-by-trial correlations were not solely driven by the main effect of target *acceleration* on TTC_eye_ and TTC_hand_, we analysed individual observers’ data using a linear model. In 13 out of the 16 observers, TTC_eye_ explained a significant portion of the variance in TTC_hand_. For example, observer S15’s TTC_hand_ performance was well predicted by TTC_eye_ (resulting in a slope of .74; *F*(2,167) = 72.27; *p* < .001; **Fig. 6B**). In the other three observers, TTC_hand_ data were not predicted well by TTC_eye_ after factoring out the main effect of target acceleration (e.g., observer S14: slope = .16*; F*(2,173) = 2.33; *p* = .13; **Fig. 6C**).

**Figure 6.**
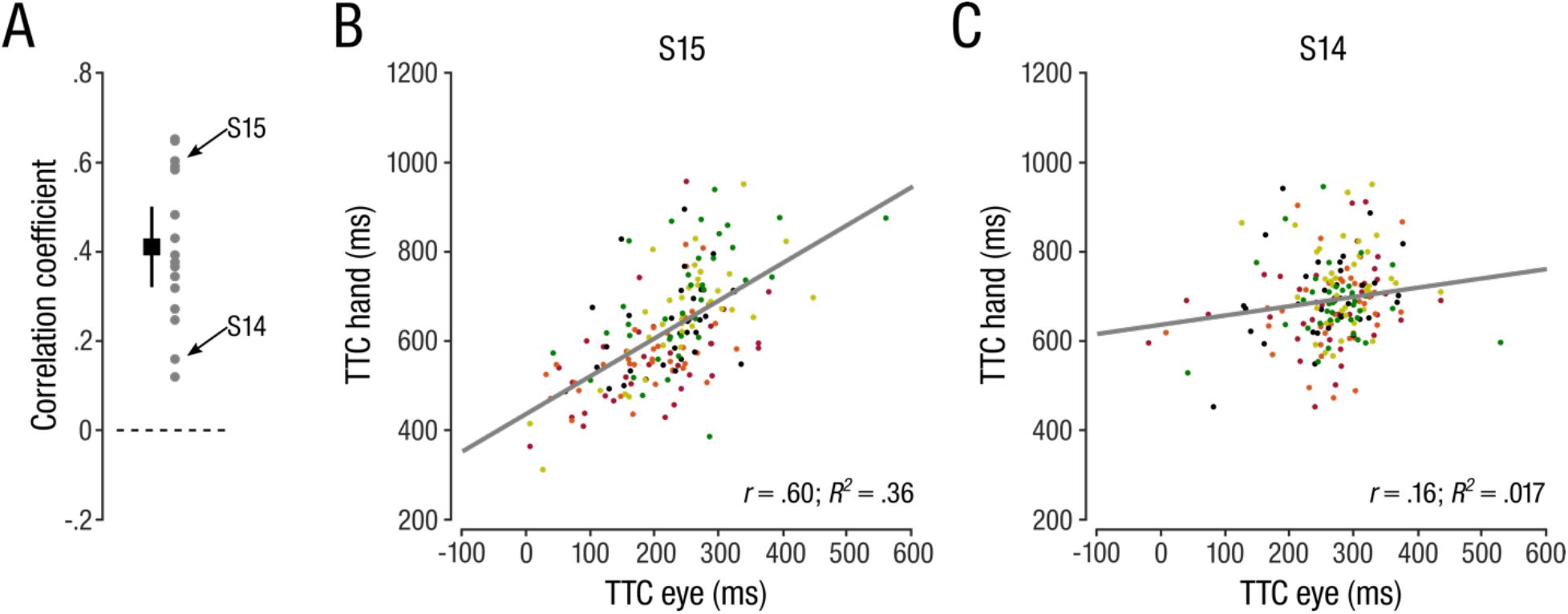
Trial-by-trial correlation between TTC_eye_ and TTC_hand_. (**A**) Mean (black square) and distribution of correlation coefficients (grey dots show individual correlation coefficients). Errorbars denote the 95% CI. (**B, C**) Scatterplot and trend lines of the trial-by-trial correlation for an observer with a weak (**B**) and for an observer with a strong (**C**) correlation between TTC_eye_ and TTC_hand_. Coloured dots represent individual trials (same convention as in Fig. 3).

### Restricting Initial Target Presentation Affects Ability to Track but not to Predict Accelerating Targets

Thus far, we have shown that observers can accurately track accelerating targets with smooth pursuit eye movements but fail to take target acceleration into account to predict target TTC. Instead, our data show that observers rely on a target velocity sample shortly before occlusion to extrapolate target motion and predict TTC. To probe this potential difference between smooth pursuit and ocular and manual interception, we next compared our results to the tracking and interception data observed when restricting the target presentation duration to either 200 or 500 ms. We expected that restricting target presentation duration would decrease an observer’s ability to track the target with smooth pursuit eye movements (e.g., Fooken et al., 2016; Kreyenmeier et al., 2017), resulting in lower sensitivity to target acceleration in visually-guided pursuit. Following the predictions of the final velocity model, we would not expect to see an impact of presentation duration on TTC prediction in eye or hand. As expected, pursuit velocity scaled with presentation duration and was significantly lower for targets that were visible for only 200 or 500 ms, compared to 800 ms (**Fig. 7A**). This observation was confirmed by a 5 (*acceleration*) x 3 (*presentation duration*) rmANOVA on average pursuit velocity, yielding a main effect of *presentation duration* (*F*(2,30) = 182.74; *p* < .001; *η_p_^2^* = .92). There was also a main effect of target *acceleration* on the average pursuit velocity (*F*(4,60) = 35.79; *p* < .001; *η_p_^2^* = .70). However, the normalized pursuit velocities show (**Fig. 7B**) show that pursuit velocity only differentiated between the different target trajectories when the targets were presented for 500 or 800 ms but not for 200 ms, which was supported by a significant *acceleration* × *presentation duration* interaction (*F*(8,120) = 19.35; *p* < .001; *η_p_^2^* = .56).

**Figure 7.**
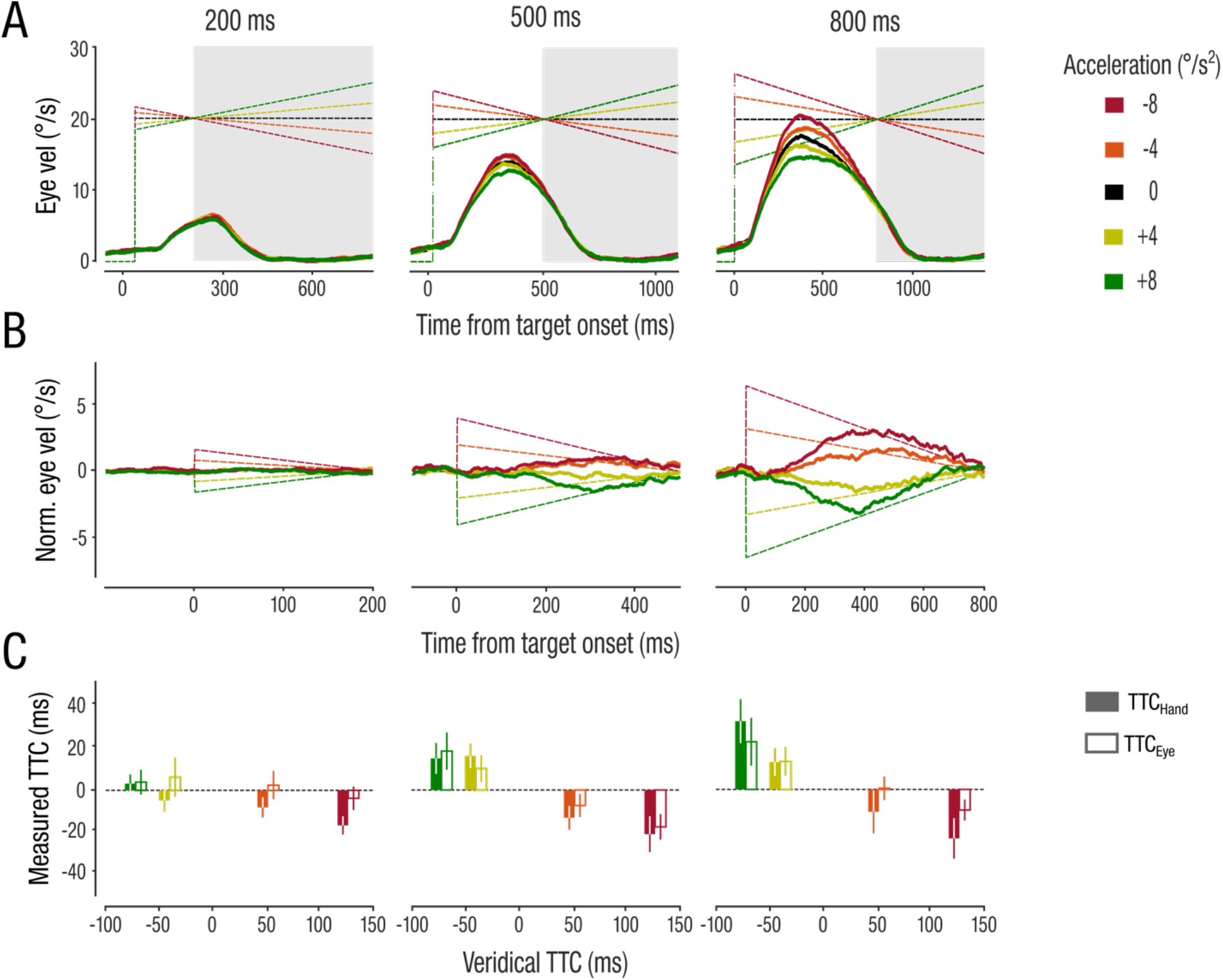
Effect of presentation time on smooth pursuit velocity and TTC. (**A**) Average de-saccaded smooth pursuit velocities. Grey zones indicate occlusion period and dashed lines represent target velocities. (**B**) Normalized pursuit velocity. (**C**) TTC estimation for eye and hand.

Whereas the smooth pursuit response scaled with presentation duration, TTC prediction was less affected (**Fig. 7C**). We ran two separate 4 (*acceleration*) x 3 (*presentation duration*) rmANOVA on normalized TTC_eye_ and TTC_hand_. For both measures, only the main effects of *acceleration* was significant (TTC_eye_: *F*(3,45) = 7.55; *p* = .007; *η_p_^2^* = .33; TTC_hand_: *F*(6,90) = 13.4; *p* < .001; *η_p_^2^* = .47), whereas neither the main effects of presentation duration, nor the interaction terms reached significance (all *F* < 2.57; *p* > .08; *η_p_^2^* < .14). These results confirm the differential impact of acceleration signals on tracking (visually-guided pursuit) and predicting (interceptive eye and hand movements). Finally, we also compared TTC_eye_ and TTC_hand_ on a trial-by-trial basis and observed—similar to the 800 ms condition—that ocular and manual interceptions were moderately correlated in both the 200 ms (*mean r* = .33 ± .17; *t*(15) = 7.0; *p* < .001; *d* = 2.0) and the 500 ms (*mean r* = .31 ± .20; *t*(15) = 6.25; *p* < .001; *d* = 1.56) conditions.

## Discussion

The current study investigates how humans integrate visual motion information to track and predict accelerating objects for manual interception. Our task required observers to track an accelerating target before a temporary occlusion and to predict the time of target reappearance by making an interceptive eye and hand movement. Our findings show that visually-guided smooth pursuit eye movements before occlusion reflected different rates of target acceleration. This suggests that acceleration information was accurately read out by the pursuit system and could potentially inform predictive eye and hand movements. However, we found that target motion prediction did not scale with acceleration when intercepting the reappearing object with either the eyes or the hand. Instead, estimates of TTC were best described by a model that relied on the final velocity of the target just before occlusion, indicating that observers based their prediction on the memory of the last available velocity signal (first-order motion; Benguigui et al., 2003; Benguigui & Bennett, 2010). This dissociation implies that vision-based tracking and prediction-based interception relied on different motion integration mechanisms of velocity and acceleration signals. These results have important implications for our understanding of how the visual and motor systems integrate (or fail to integrate) non-linear motion signals when interacting with accelerating objects.

### Dissociation between Tracking and Predicting

To successfully interact with a moving object, e.g., to catch or intercept it, we must continuously monitor its dynamically changing motion trajectory. When we lose sight of the object, e.g., due to a temporarily occlusion, we need to quickly form a prediction of current and future object motion. Naturally-moving objects do not necessarily move at constant velocity but can suddenly accelerate or decelerate. Forming a predictive model that can capture dynamically changing object motion is therefore an integral part of everyday actions.

The abilities to track and predict the motion trajectory of objects that move at constant velocity are often closely linked (Makin & Poliakoff, 2011). Accurate tracking of a moving object with smooth pursuit eye movements can enhance temporal (Bennett et al., 2010) and spatial (Spering et al., 2011) predictions of target trajectories. Our results suggest that the close link between tracking and predicting object motion does not extend to accelerating targets. Instead, we found a dissociation between the ability to track and predict accelerating objects when aiming to intercept them. Specifically, we show that observers’ eye movements systematically reflected acceleration information for target presentations of more than 200 ms. However, irrespective of how long observers had time to track and potentially integrate acceleration signals, the timing of manual and ocular interception indicated inability to consider acceleration in our task.

Our finding of different acceleration integration for tracking and interception is congruent with two sets of literature that have typically tested both behaviors—tracking and interception—separately. First, the smooth pursuit system can be sensitive to acceleration signals when probing it with velocity perturbations (Brostek et al., 2017; Tavassoli & Ringach 2010). Moreover, predictive pursuit during a target’s occlusion period scales with the target’s acceleration (Bennett & Barnes, 2006; Bennett et al., 2007; Bennett & Benguigui, 2013). These findings suggest that the oculomotor system can extract acceleration signals even to predictively drive pursuit. This capability to decode and utilize acceleration signals in the pursuit system is surprising, considering that perceptual judgements of target prediction are largely insensitive to acceleration and best predicted by first-order motion extrapolation (Bennett & Benguigui, 2013; Bennguigui et al., 2003; Benguigui & Bennett, 2010). Second, interceptive hand movements are comparatively unresponsive to visual acceleration, reflected in systematic errors when intercepting accelerating targets (Benguigui et al., 2003; Brenner & Smeets, 2015; Brenner et al., 2016; Dubrowski & Carnahan, 2002; Port et al., 1997). These systematic interception errors can be explained by a failure of observers to predict accelerating target motion overcome sensorimotor delays (Brenner & Smeets, 2015). Here we extend these previous findings by showing that observers also fail to consider acceleration in their prediction of target motion to overcome temporary target occlusion.

One possible explanation for the observed dissociation between tracking and predicting accelerating targets could be that visually-guided tracking can rely on detecting and updating the changing target velocity over time and might thus not require a direct consideration of the acceleration signal. Conversely, to intercept accelerating targets, observers need to consider an explicit readout of target acceleration to form a prediction of target motion to overcome sensorimotor delays (Brenner & Smeets, 1997; 2015) or a temporary occlusion of the moving object. The finding that acceleration is not used during manual interception suggests that observers continuously update their judgment of target velocity but cannot integrating acceleration signals to inform their prediction. Instead, they used a single velocity sample just before target occlusion to predict TTC. Our analyses revealed that velocity samples of ~100 ms and ~60 ms before occlusion onset best predicted the timing of manual interception (TTC_hand_) and the landing time of the targeting saccades (TTC_eye_), respectively. Thus, TTC_eye_ and TTC_hand_ relied on very similar velocity samples, suggesting that both predictive eye and hand movements were tightly coupled and similarly insensitive to acceleration.

### Extending the Eye-Hand Link to Prediction-Based Actions

We observed trial-by-trial correlations between TTC_eye_ and TTC_hand_, extending the known close coupling of eye and hand movements during visually-guided actions (Hayhoe, 2017; De Brouwer et al., 2021) to predictive actions (Binaee & Diaz, 2019). During visually-guided reaching, observers commonly shift their eyes to the reach target prior to hand movement execution (Ballard et al., 1992; Johansson et al. 2001; Neggers & Bekkering, 2000; Horstmann & Hoffmann, 2005; Barany et al., 2020; Land & Hayhoe, 2001). When intercepting moving targets, observers naturally track the target with their eyes, even when no explicit instruction to do so is given (Mrotek & Soechting, 2007). In interception tasks, eye and hand movement endpoints are also correlated (Kreyenmeier et al., 2017; Li et al., 2018; Fooken et al., 2021). Our results extend these findings in two ways. First, correlations of predictive interception accuracies in eye and hand reveal that the coordinated control of eye and hand movements also applies to memory-based actions. Second, correlations of temporally-based estimations show that eye and hand movements can be correlated not just in the spatial, but also in the temporal domain. Although the eyes reach the interception location several hundred milliseconds before the hand, the timing of interceptive eye and hand movements were correlated on a trial-by-trial basis in ~80% of the observers. A strong eye-hand link is expected when intercepting targets that move unpredictably and are partially occluded from view (Fooken et al., 2021). If acceleration is indeed not considered in a predictive model of target motion, the motion of accelerating targets becomes highly unpredictable, and observers rely on their eye movements to continuously update their prediction of the target motion (Brenner & Smeets, 2018; de la Malla et al., 2019).

### Assessing Model Predictions of Accelerating Motion Integration

Given the limited perceptual sensitivity to acceleration, and the lack of acceleration tuning in key motion-sensitive cortical areas (Lisberger & Movshon, 1999; Price et al., 2005) the question arises what information observers rely on when interacting with accelerating objects in everyday life. It is well known that humans use physical laws of motion, such as gravity, which are learned throughout the lifespan, as an implicit prior when interacting with real-world objects (Jörges & López-Moliner, 2017). For instance, observers are more accurate when tracking and predicting simulated fly balls that move with natural gravity compared to balls that do not (0g), or that are unnaturally impacted by gravity (2g; Delle Monache et al., 2019; Bosco et al., 2012). Moreover, humans are generally able to quickly adapt movements after only a few trial repetitions (Ruttle et al., 2021). For example, improvements in the ability to manually intercept (Brenner et al., 2016) and predictively pursue (Bennett & Barnes, 2006) accelerating targets after a few (eight to twelve) repetitions of the same acceleration rate have been reported. These findings suggest that observers might be able to form short-term and long-term (naturalistic) priors to counteract the lack acceleration signal integration.

In conclusion, our study shows that the extraction of acceleration signals during smooth pursuit was not used for an acceleration-based prediction of the target’s motion to inform manual interception. Instead, the timing of predictive eye movements and manual interception were best predicted by an extrapolation of target motion based on a velocity sample shortly before target occlusion. Our results suggest that acceleration signals are differently integrated for visually-guided tracking and prediction-guided interception. Both interceptive eye and hand movements showed strikingly similar insensitivity to acceleration and were correlated on a trial-by-trial basis, indicating a strong coupling between both effectors during prediction-guided interception tasks.

## Materials and Methods

### Observers

We tested 16 human adults (seven females; mean age 26.8 ± 5.1 years, range 19-37 years; including authors PK and LK) in this study. The sample size was determined using a priori power analyses in two steps (G*Power version 3.1; Faul et al., 2009): First, based on effect sized reported in Brenner and colleagues (2016), we determined that a sample size of *n* = 5 was required to detect systematic interception errors caused by target acceleration (power = .80; alpha = .05). Second, to ensure that our sample size also provided sufficient power to detect an effect of acceleration on eye movements, we used the regression slopes reported previously (e.g., Bennett et al., 2007), yielding our final sample size of *n* = 16 (power = .80; alpha = .05). All participants had normal or corrected-to-normal visual acuity and reported no history of neurological, psychiatric, or eye disease. The study protocol was carried out in accordance with the Declaration of Helsinki and was approved by the Behavioural Research Ethics Board at the University of British Columbia. Participants gave written informed consent before participating and were compensated at the rate of $10/h.

### Apparatus

Participants performed the task in a dimly lit laboratory, viewing the stimuli binocularly at a distance of 44 cm. A PROPixx video-projector with a resolution of 1,280 × 1024 pixels and a refresh rate of 120 Hz (VPixx Technologies, Saint-Bruno, QC, Canada) was used to back-project the stimuli onto a 41.8 × 33.4 cm translucent screen. The position of participants’ right eye was recorded using a video-based eye tracker (Eyelink 1000 Tower Mount, SR Research, Ottawa, ON, Canada) with a sampling rate of 1 kHz. A combined chin and forehead rest minimized head movements during the experiment. A small magnetic sensor was attached to the tip of participants’ right index finger to record their 3D hand movements with a 3D Guidance trakSTAR (Ascension Technology, Shelburne, VT) at a sampling rate of 120 Hz. The experiment was programmed in Matlab (MathWorks, Natick, MA, USA), using the Psychophysics (version 3.0.16; Kleiner et al., 2007) and EyeLink toolboxes (Cornelissen et al., 2002). Stimulus display and data collection were controlled by a PC (graphics card: NVIDIA GeForce GTX 1060).

### Stimuli and Procedure

Participants viewed and intercepted a small black disc (0.35° in diameter; 6.23 cd/m^2^) that moved across a light grey background (229.8 cd/m^2^) and then disappeared behind an occluder, a grey (181.3 cd/m^2^) bar, that extended 13.4° from the horizontal midline into the right half of the screen. Each trial started with the disc shown on the left side of the screen. Participants had to fixate the target (400-800 ms) and place their index finger on a designated start position on the table. Upon successful fixation, the target started moving to the right, either with a fixed velocity (0°/s^2^) or constantly accelerating at different rates (−8, −4, 4, 8 °/s^2^). The target was shown for 200, 500, or 800 ms before occlusion. Participants were instructed to follow the disc closely with their eyes during the initial presentation and to manually intercept the target at the time they expected it to reappear behind the occluder. Target presentation ended either with the time of interception, or 100 ms after target reappearance. Initial target position and velocity were matched for each presentation time and acceleration so that all targets reached the same position and velocity at the time of occlusion. The time of target reappearance (equivalent to time-to-contact, TTC) depended on the target’s acceleration rate. Therefore, successful interception required adjusting the timing of manual interception to target acceleration. Observers were instructed to intercept as closely to the border of the occluder as possible at the time they expected the target to reappear. Feedback about the interception performance was provided at the time of interception by showing a red and a black dot, indicating the interception position and the actual target location at the time of interception, respectively. The combination of acceleration and presentation time resulted in 15 experimental conditions, presented in random order within each block of trials. Each participant completed 40 trials per condition, resulting in 600 trials total, presented in eight blocks of 75 trials each. The experiment took approximately 90 minutes.

### Eye and Hand Movement Recordings and Analyses

The data were pre-processed offline using custom-made routines in MATLAB. Eye velocity and acceleration were calculated as the first and second derivatives of the eye position signals over time. Position and velocity profiles were filtered using a low-pass, second-order Butterworth filter with cut-off frequencies of 15 Hz (position) and 30 Hz (velocity). Saccades were detected when five consecutive frames exceeded a fixed velocity criterion of 30°/s; saccade on- and offsets were then determined as the nearest reversal in the sign of acceleration. For the analyses of the de-saccaded smooth pursuit eye movements, the identified saccades ± 25 ms were removed from pursuit velocity traces and replaced by linear extrapolation between the last velocity before saccade onset and the first velocity sample after saccade offset. Pursuit onset in de-saccaded traces was detected within a 300-ms interval around stimulus motion onset (starting 100 ms before onset) in each individual trace. We first fitted each 2D position trace with a piecewise linear function, consisting of two linear segments and one breakpoint. The least-squares fitting error was then minimized iteratively (using the function lsqnonlin in MATLAB) to identify the best location of the breakpoint, defined as the time of pursuit onset.

The magnetic hand tracker recorded the 2D screen-centred interception position as well as the participant’s hand movement in 3D space. Hand position data were up-sampled to 1 kHz by linear interpolation for precise temporal comparison with eye movement data. Hand latency was computed offline as the first sample exceeding a velocity threshold of 5 cm/s following stimulus onset. Hand movement offset was detected when the finger intercepted the screen (within ± 0.16 cm of the screen).

All trials were manually inspected. We excluded trials with blinks during the task and trials in which the eye tracker lost the signal (3.0% of trials across participants). Trials were also excluded when participants tried to intercept the target on top of the occluder (2.6% of trials), or when no interception was detected within 600 ms of target reappearance (3.4% of trials).

### Data Analyses

The primary aim of the current study was to compare observers’ ability to track and intercept accelerating targets. We assessed observers’ ability to accurately track accelerating targets with smooth pursuit eye movements during the initial target presentation (visually-guided smooth pursuit). Because observers stereotypically made an initial catch-up saccade to align the eyes with the target (de Brouwer et al., 2002), we restricted our analyses to the closed-loop pursuit phase after this initial catch-up saccade. If an initial catchup saccade was followed by second catchup saccade within 50 ms, we used the second catchup saccade offset as the beginning of the smooth tracking phase. We then measured the average eye acceleration over time between the offset of the initial catchup saccade and the onset of target occlusion. Because of the relatively noisy signal of eye acceleration, we additionally compared the smooth pursuit velocities at two critical time points: (1) The *initial pursuit velocity* was computed from a 50 ms time window following the initial catch-up saccade and (2) the *final pursuit velocity* was measured during the last 50 ms before target occlusion onset. Pursuit gain was typically less than 1 and observers showed anticipatory slowing in the pursuit before target occlusion. To account for these general biases, we normalized pursuit velocity and acceleration measures by subtracting the control condition (0°/s^2^) from the experimental conditions. Following target occlusion, observers typically stopped pursuing the target with smooth pursuit eye movements and used predictive saccades to bring the eyes to the interception location. The primary predictive saccade was determined as the saccade that brought the eye within 3.5° from the occluder’s right edge (to allow for systematic undershooting of saccades). The landing time of the eye (TTC_eye_) was determined as the offset time of the predictive saccade. If additional saccades were made to correct the undershooting, we used the offset time of the last corrective saccade that was initiated from within the occluder. To assess whether target acceleration was taken into account in manual interception, we calculated the constant interception error as the difference between the time of interception and the veridical time the target would have been at the interception location. We also analyzed hand movement latency, peak velocity, movement time, and the interception time (TTC_hand_).

#### Model comparison

The eyes arrived at the interception location several hundred milliseconds before the time of manual interception. Thus, to compare biases between TTC_eye_ and TTC_hand_ we normalized TTC_eye_ and TTC_hand_ by subtracting the control condition (0°/s^2^) from the experimental conditions. We then compared three different models on how observers might have predicted TTC. We calculated targets’ TTC based on these models (**Fig. 5A-C**): For the average velocity model, TTC was calculated as *TTC(x) = d* / vavg(x), for the final velocity model, TTC was calculated as *TTC*(*x*) = *d* / v_final_(*x*), and for the acceleration model as 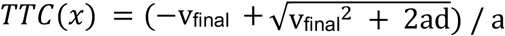. Here, *x* indicates the 5 different target trajectories, *d* the distance of the occluder, *v_avg_* the average velocity of the target during the initial presentation duration, *v_final_* the final target velocity at the time of occlusion onset, and *a* the rate of target acceleration. We then evaluated the fit of the different model predictions of TTC to our observed TTC_eye_ and TTC_hand_ data by calculating the root-mean-square error (RMSE) for each observer. To further analyse which target velocity sample best described each observers’ ocular and manual TTC biases, we predicted TTC based on each timepoint along each target’s velocity trajectory and determined the timepoint that produced the smallest RMSE, separately for TTC_eye_ and TTC_hand_. We coined this timepoint the eye and hand prediction times, respectively.

### Statistical Analyses

For all outcome variables, we calculated the median across trials for each individual observer. To assess differences in our experimental conditions, we ran repeated-measures analyses of variance (rmANOVA) and post-hoc t-tests. Violation of sphericity was assessed using the Mauchly’s test and p-values were Greenhouse-Geisser corrected in case of significance. Bonforroni corrections were used for any post-hoc comparisons. In order to investigate whether TTC_eye_ predicted TTC_hand_ on a trial-by-trial basis, we calculated Pearson’s correlation for each observer individually and confirmed our findings using a linear model that additionally contained the acceleration condition as a categorial predictor. A full linear model (containing main and interaction effects of *TTC_eye_* and target *acceleration*):

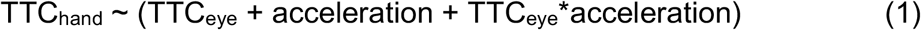

was compared to a partial linear model that only contained a main effect of target *acceleration* and the interaction term to obtain the main effect of TTC_eye_ on TTC_hand_:

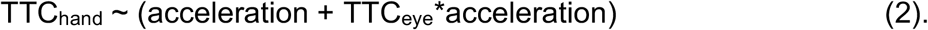

All statistical analyses were performed in R (version 3.3.2; R Core Team) with an alpha level of .05.

## Acknowledgements

This work was supported by a UBC International Doctoral Fellowship to PK, a Deutsche Forschungsgemeinschaft (DFG) Research Fellowships to JF (grant FO 1347/1-1), and an NSERC Discovery Grant and Accelerator Supplement to MS. The authors thank members of the Spering lab for comments on an earlier version of the manuscript.

## Author Note

The study design and parts of the analysis were preregistered (https://osf.io/adq9v), and data will be made available on the Open Science Framework. The model comparison was not included in the preregistration. Preliminary data were presented at the 2021 virtual meeting of the Vision Sciences Society (Kreyenmeier et al., 2021).

